# DNA methylation age studies of humpback whales

**DOI:** 10.1101/2022.08.15.503952

**Authors:** Steve Horvath, Amin Haghani, Joseph A. Zoller, Zhe Fei, Martine Bérubé, Jooke Robbins

**Author notes:** Joint First Author (Steve Horvath, Amin Haghani).

## Abstract

Several previous studies have described epigenetic clocks for whale and dolphin species. Here we present a novel and highly robust epigenetic clock for the humpback whale based on methylation levels measured using the Mammalian Methylation Array platform. Skin samples were obtained from 76 individuals that had been studied since their birth year and known to range in age from <1 to 39.5 years. The humpback whale clock provided a highly accurate estimate of chronological age (R=0.96, median error 2.2 years) according to a leave-one-out cross validation analysis. We applied this clock to an independent set of samples from humpback whales of unknown exact age but with sighting histories that were as long as or longer than the upper 20% of the available known-age range. Although there was a strong correlation with minimum age (R=0.89), the clock underestimated age in these older animals by a median error of at least 7.8 years. Finally, we applied the humpback clock to publicly available methylation data from beluga whales. In this data set from a different species, the humpback clock provided an age correlation of R=0.78. While a DNAm age estimator has previously been described for humpback whales, this is the first such clock shown to apply to another cetacean species as well. Our humpback whale clock built from well-studied population lends itself for understanding humpback populations that otherwise lack age data.

## INTRODUCTION

Age is important for understanding fundamental aspects of biology and ecology, both at the individual and population level. However, some species, such as the baleen whales, cannot be reliably aged chronologically nor, in most cases, even assigned to all biologically important age classes. Early efforts to identify age-related markers focused on whales killed during historical commercial whaling (Chittleborough 1959, 1965). Growth layer groups in waxy ear canal deposits (ear plugs) were thought to reflect chronological age; however, there is two-fold uncertainty in ear plug based estimates of age and lifespan, depending on assumptions about layer deposition (Gabriele et al. 2010; Best 2011). Aspartic acid racemization of the eye lens has also been used to age baleen whales (Garde et al. 2007; Rosa et al. 2013; Boye et al. 2020). However, as in the case of ear plugs, the necessary samples can only be obtained from dead animals. Such aging techniques are therefore not well-suited for the study of long-lived species of conservation concern.

The only precise data on chronological age in baleen whales comes from the few long-term studies of living individuals uniquely identified in their year of birth. (Dunshea et al. 2011; Olsen et al. 2012; Jarman et al. 2015) That research has demonstrated significant relationships between age class and behavior, association patterns, distribution, reproduction, survival and risk of impacts from human activities (e.g., Cubbage & Calambokidis 1987; Clapham 1992, 1994; Ramp et al. 2010; Robbins et al. 2015; Knowlton et al. 2016). As such, the development of reliable markers of chronological age continues to be highly desirable for scientific and conservation applications. Small samples of skin and shallow blubber tissue can be collected from living whales of known age, and these been used to evaluate the potential of aging methods based on fatty acids signatures (Herman et al. 2009), telomers (Dunshea et al. 2011; Olsen et al. 2012) and, more recently, epigenetic markers (Polanowski et al. 2014).

Previous studies have shown that telomere length does not accurately predict chronological age in humpback whales and vertebrates (Dunshea et al. 2011; Olsen et al. 2012; Jarman et al. 2015). By contrast, there is strong evidence that epigenetic age estimators based on DNA methylation data lend themselves for developing highly accurate estimators of chronological age in humans and other mammals (Jarman et al. 2015; Jylhava et al. 2017; Horvath & Raj 2018). Specific cytosine-guanine dinucleotides (CpGs) can be methylated to generate 5-methylcytosine, a chemical modification that may affects gene expression (Razin & Cedar 1991; Field et al. 2018).

Several DNA methylation-based age predictors, referred to as epigenetic clocks, have been developed for humans, mice and other mammals (Bocklandt et al. 2011; Hannum et al. 2013; Horvath 2013). Epigenetic clocks have also been developed for several cetacean species, including minke whales, *Balaenoptera acutorostrata* (Tanabe et al. 2020), humpback whales, *Megaptera novaeangliae* (Polanowski et al. 2014), bottlenose dolphins, *Tursiops truncatus* (Beal et al. 2019; Barratclough et al. 2021; Robeck et al. 2021), beluga whales, *Delphinapterus leucas* (Bors et al. 2020), and more generally odontocetes (Robeck et al. 2021).

Here, we present a significant advance in the aging of humpback whales based on DNA methylation data. This epigenetic clock leverages data and skin samples from a long-term study of North Atlantic humpback whales in the Gulf of Maine and a robust measurement platform (the Mammalian Methylation Array). We applied the resulting humpback whale clock to samples from humpback whales whose birth year was unknown but in most cases preceded individuals in the training data. Finally, recognizing that age data are rarely available for cetacean species, we show that the humpback clock appears to be informative for skin samples from another cetacean species, the beluga whale.

## Results

We obtained methylation profiles of DNA from skin of humpback whales (n=91, Table 1) using the mammalian methylation array (Methods).

**Table 1.** Description of the skin methylation data from humpback whales. N=Total number of samples per data set. The training data involved animals whose ages were known because they were studied since their year of birth. The test data included animals of unknown exact age with sighting histories that were generally as long as or longer than the upper 20% of the available known-age range. Number of females. Age: mean, minimum and maximum.

| Data | Confidence | N | No. Female | Mean Age | Min. Age | Max. Age |
| --- | --- | --- | --- | --- | --- | --- |
| Training | Chronological age known | 76 | 39 | 16.9 | 0.4 | 39.5 |
| Test | Minimum age only | 14 | 11 | 35.5+ | 13.3+ | 43.7+ |

### Epigenetic clocks

We developed the humpback whale clock based on a subset of 76 skin samples from individuals known to range in age from <1 to 39.5 years. This clock was then applied to a second set of samples for which only the minimum age was known. One of these minimum ages was in the primary range of the training data set while the minimum possible age of the rest averaged 37.05 years.

To arrive at unbiased estimates of the epigenetic clocks, we performed cross-validation analyses in the training data and obtained estimates of the age correlation R (defined as Pearson correlation between the age estimate, DNAm age and chronological age), as well as the median absolute error. The humpback clock provide a highly accurate estimate of chronological age (R=0.96, median error 2.2 years, **Figure 1A**) according to a leave one out cross validation analysis in the training data. When applied to samples from humpbacks for whom only a lower bound of the chronological age was available (**Figure 1B**), the clock underestimated the minimum age by a median error of at least 7.8 years. The magnitude of age under-estimation is not known for these samples, but there was a strong correlation with minimum age (R=0.89). Finally, we applied the humpback clock to publicly available skin methylation data from beluga whales which were profiled on the same array platform (Bors et al. 2020). In this test data set from this different species, the humpback whale clock achieved an age correlation of R=0.78 and a median error of 2.8 years (**Figure 1C**).

**Figure 1.**
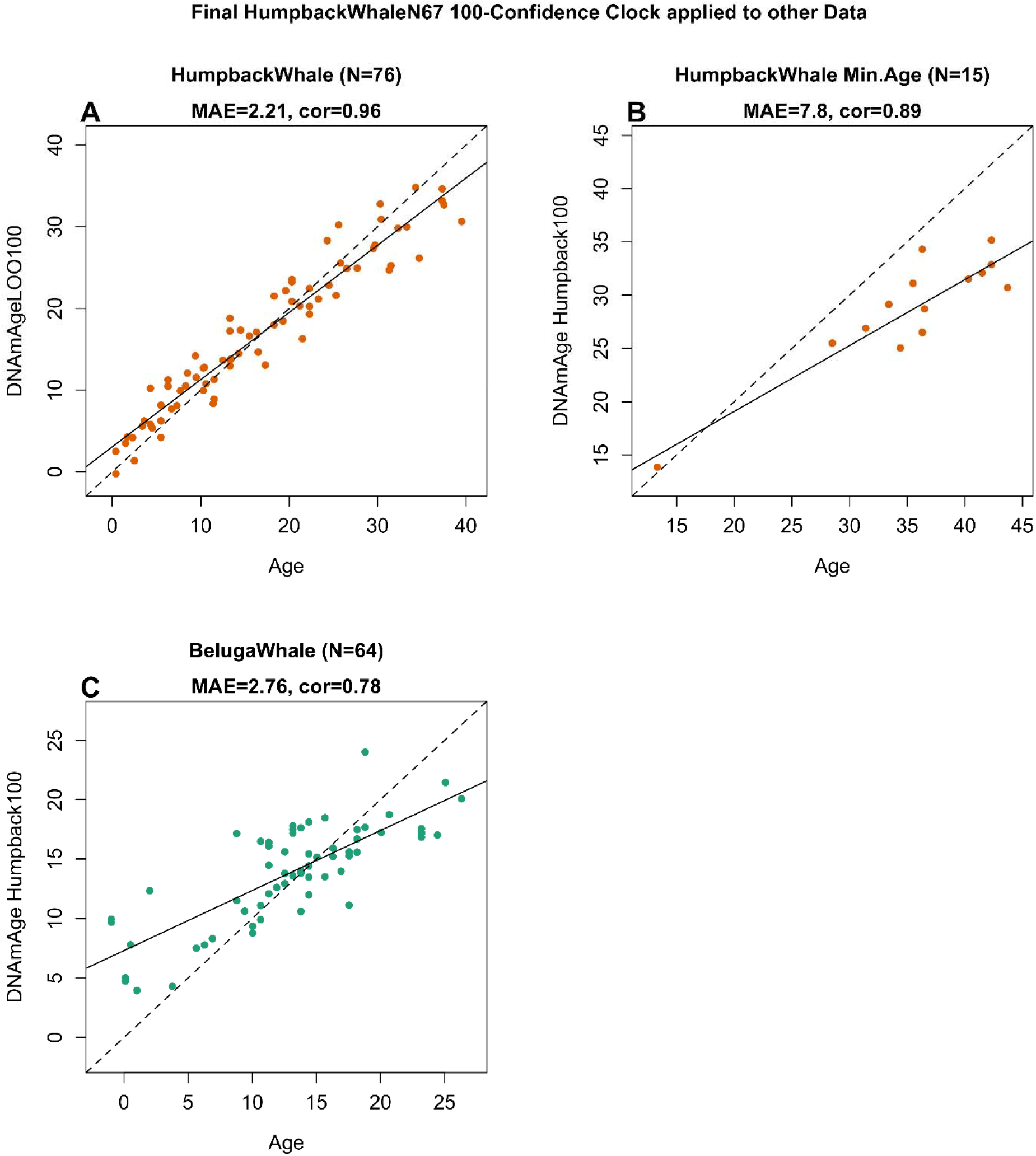
Humpback whale clock evaluated in different data sets. A) Chronological age versus a leave one out (LOO) cross validation estimates of DNA methylation age (y-axis, in units of years). These data were comprised of n=76 animals whose chronological ages were known (100% confidence). B) Independent test set comprised of 15 skin samples from humpback whales of unknown exact age but with sighting histories that were generally as long as or longer than the upper 20% of the available known-age range. C) Test set comprised of skin samples from beluga whales. Each panel reports the sample size, correlation coefficient, median absolute error (MAE) in units of years.

### Epigenome-wide association study (EWAS) of age in humpback whales

The Mammalian Methylation Array is designed based on the conserved stretches of DNA in all mammals (Arneson et al. 2022). Thus, even though an adequate humpback genome assembly was not available at time, the CpG level analysis could be based on killer whale annotations, which contained 30,467 probes adjacent to 5,730 genes throughout the genome. A subset of 2,451 CpGs were related to age in humpback whale skin (p<10^-6^, **Figure 2A**). Some of the top age-related changes in humpback whale included hypermethylation in *LOC117197173* downstream, *NTM* intron, *PRUNE2* intron, *TTN* intron, and hypermethylation in *LOC101287766* upstream, *EN1* promoter, *NOL4* exon, and *TWIST1* downstream (**Figure 2D**). The strong positive and negative correlations were quite strong for these sites (r>=0.89 and r<-0.9, respectively). Our EWAS of age in humpback whales implicated CpGs that were located near developmental genes and targets of polycomb repressor complex 2 (**Supplementary Figure 2**) similar to what has been seen in other mammalian species. We observed both age related gain and loss of methylation in genic and intergenic regions. But CpGs located in gene promoters and 5’UTRs mainly gained methylation with age (**Figure 2B**). Similarly, CpGs in CpG islands, which are located near promoters, showed age related gain of methylation (**Figure 2C**). Collectively, these results mirror those in other mammalian species (Lu et al. 2021).

**Figure 2.**
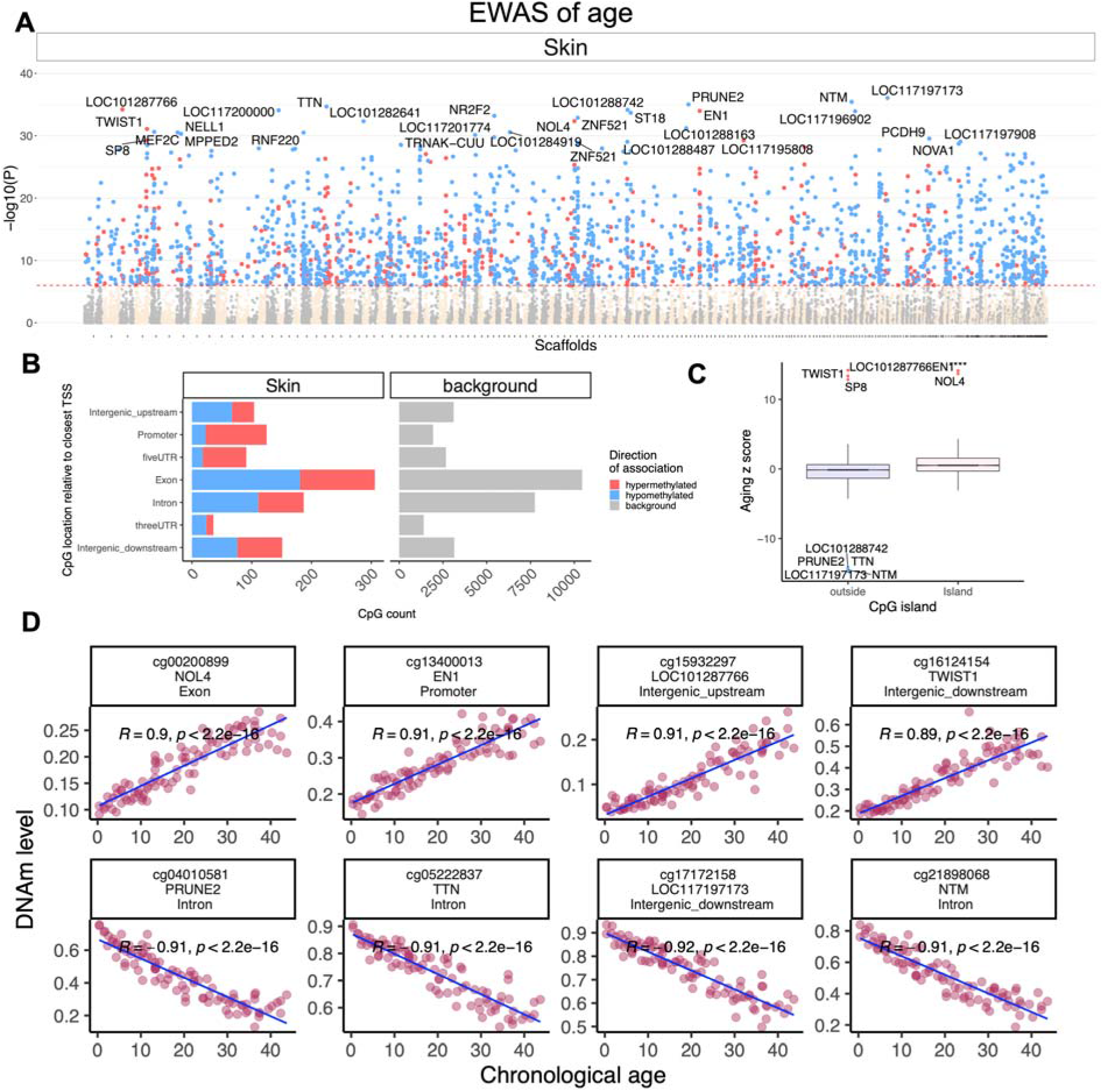
Epigenome-wide association (EWAS) of chronological age in humpback whale skin. A) Manhattan plots of the EWAS of chronological age. We used the n=91 available skin samples. Since the humpback whale genome assembly was not available, the coordinates were estimated based on the alignment of Mammalian array probes to Killer_whale.Orcinus_orca.GCF_000331955.2_Oorc_1.1 genome assembly. The direction of associations with p < 10^-6^ (red dotted line) is highlighted by red (hypermethylated) and blue (hypomethylated) colors. The top 30 CpGs were labeled by the neighboring genes. B) The location of top CpGs relative to the adjacent transcriptional start site. The grey color in the last panel represents the location of 3,0467 Mammalian Methylation Array probes mapped to the killer whale genome. C) Box plot analysis of DNAm aging association by CpG island status. The top age related CpGs in each tissue are labeled by adjacent genes. **** p<10^-4^. D) Scatter plots of the top age related CpGs in humpback whale skin.

#### Sex-specific DNA methylation patterns

Our study consisted of a balanced distribution of samples from both sexes across the validated age range. This allowed us to examine sex differences in the humpback whale. A total of 833 CpGs were differentially methylated by sex in the humpback whale skin at a significance level p<10^-6^ (Female vs Males: 407 hypomethylated, 426 hypermethylated **Figure 3A**). We could not directly evaluate whether these CpGs are located on sex chromosomes in the humpback whale. But the majority (n=805) of these CpGs were located on the *human* X chromosome, which indicates that the identified whale scaffolds are sex chromosome homologues. The genes with sex-related differential methylation were related to presynaptic function, cognitive performance (in mice), histone regulation, protein ubiquitination, heart function, and X-linked targets (**Figure 4**).

**Figure 3.**
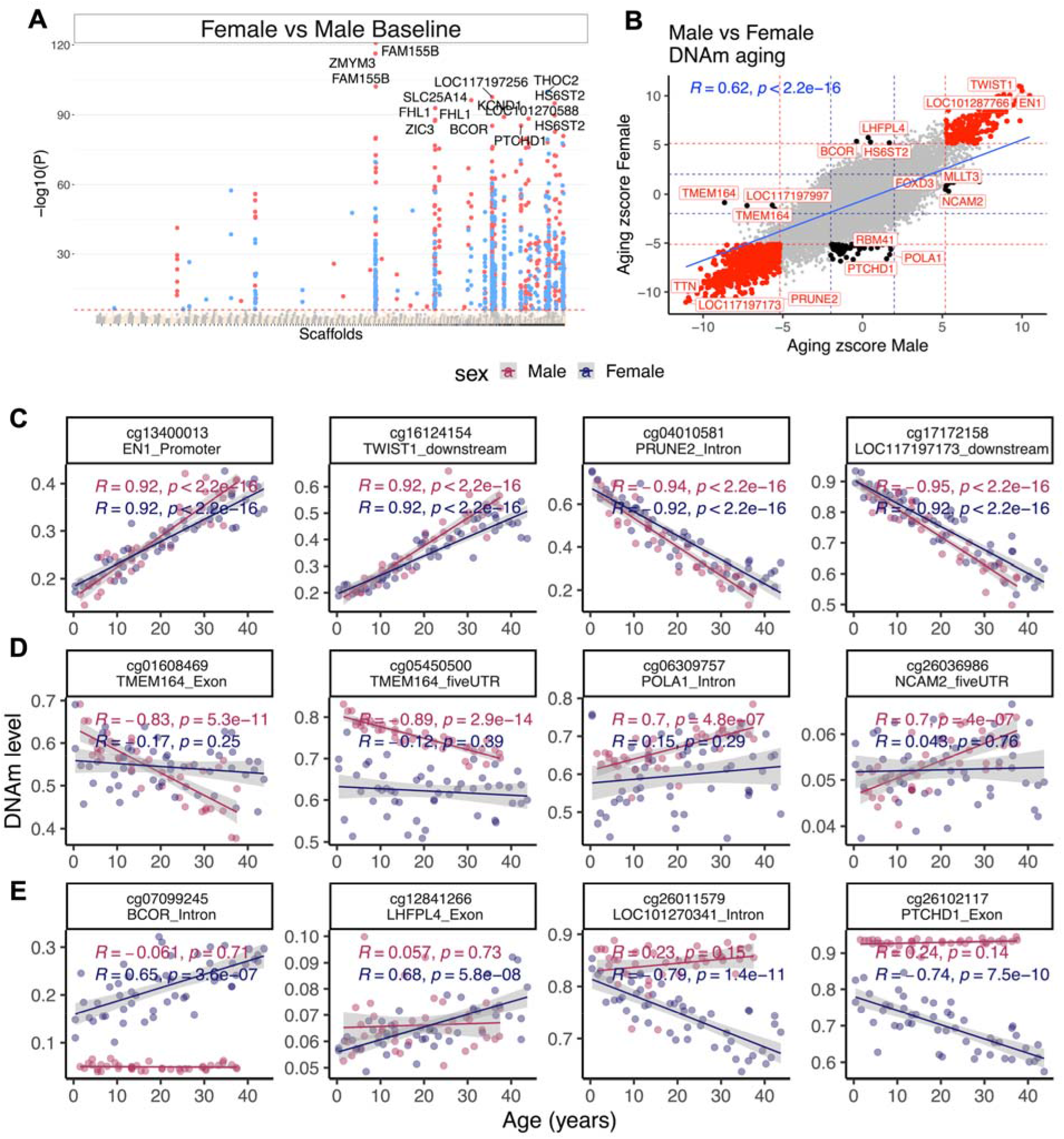
Sex alters DNAm profile of the humpback whale skin. **A)** Manhattan plot of the EWAS of sex. Co-variate: chronological age. Sample size: males, 40; Females, 50. The coordinates were estimated based on the alignment of Mammalian array probes to Killer_whale.Orcinus_orca.GCF_000331955.2_Oorc_1.1 genome assembly genome assembly. The direction of associations with p < 10^-6^ (red dotted line) is highlighted by red (hypermethylated) and blue (hypomethylated) colors. The top 15 CpGs were labeled by the neighboring genes. B) Sector plot of DNA methylation aging by sex in humpback whale. Red dotted line: p<10^-6^; blue dotted line: p>0.05; Red dots: shared CpGs; black dots: distinct changes between males and females. Scatter plots of selected CpGs that change with age in both (C), or only male (D), or female (E) humpback whales.

**Figure 4.**
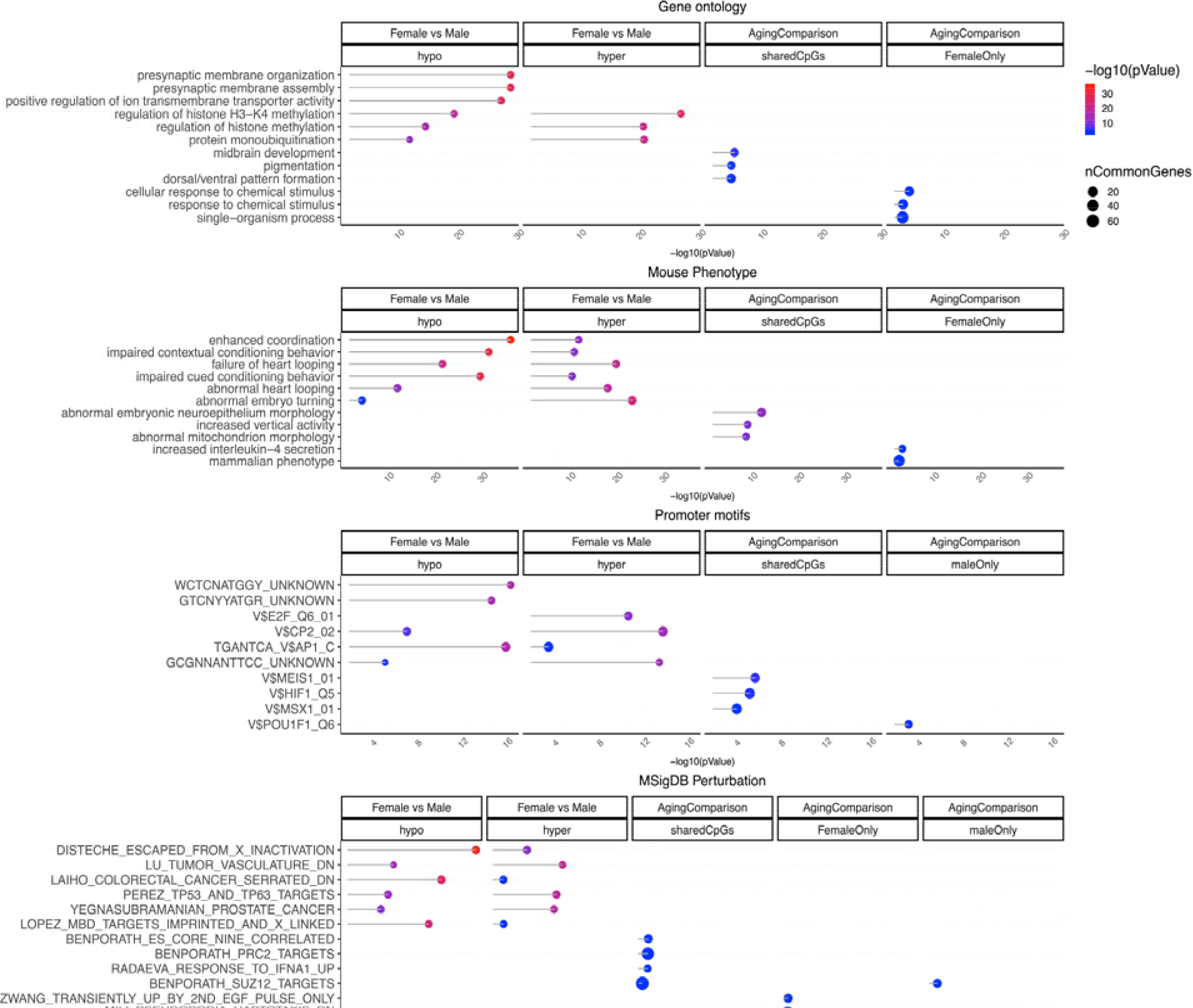
Gene set enrichment analysis of DNAm aging and sex differences in humpback whale skin. The gene level enrichment was done using GREAT analysis and human Hg19 background. Datasets: gene ontology, mouse phenotypes, promoter motifs, and MSigDB Perturbation. The results were filtered for significance at p < 10^-3^.

While aging effects on DNAm levels were highly correlated (Pearson r = 0.6) between sexes, our analysis identified a subset of 60 CpGs that only changed with age in females, and 18 CpGs that only changed in males (**Figure 3B**). For example, CpGs in the *POLA1* intron, and *NCAM2* 5’UTR only exhibited a significant age related gain in methylation in males (**Figure 3C**). By contrast, CpGs in the *LHFPL4* exon and *BCOR* intron only exhibited a significant age related gain in methylation in females (**Figure 3D**). CpGs that correlated significantly with age in female humpback whales but not male whales are located near genes related to responses to chemical stimulus and IL4 secretion (**Figure 4**).

## Discussion

Previous studies have reported on epigenetic clocks for cetacean species, including humpback whales (Polanowski et al. 2014; Beal et al. 2019; Bors et al. 2020; Tanabe et al. 2020; Barratclough et al. 2021; Robeck et al. 2021). In prior work on humpback whales (Polanowski et al. 2014), CpG sites were selected for based on their association with age in other species. An assay was then developed based on the best three of seven sites that showed promising correlations with humpback whale age. The study by Polanowski (2014) provided the first epigenetic clock in humpback whales using forty North Atlantic humpback whales (of known age) sampled in the Gulf of Maine. The present study used samples from the same population, but with roughly double the sample size and using the Mammalian Methylation Array platform that robustly measures methylation levels in highly conserved cytosines (Arneson et al. 2022). We were not able to directly compare the several epigenetic clock that lend themselves for measuring age in humpback whale due to the lack of a suitable test set. While we do not have independent data to evaluate our humpback clock (or those from other publications), the high correlation found in minimum age samples across older humpback whales (**Figure 1B**) can be explained through a fortunate sampling effect: a similar time from birth to first detection of these animals in this particular long-term population study.

Our focus on highly conserved CpGs explains why our humpback whale clock worked well in beluga whales (**Figure 1C**). We briefly mention that we have developed universal pan mammalian methylation clocks that were trained on the basis of a couple of hundred mammalian species (Lu et al. 2021).

According to our cross validation estimates, the humpback whale clock slightly overestimates the age of individuals known to be young, and to underestimate the age of those known to be old. The same “regression-to-the mean” pattern was evident in the previous independent study of humpback whales (Polanowski et al. 2014), as well as in the study of beluga whales based on the Mammalian Methylation Array platform (Bors et al. 2020). To counter this regression-to-the mean effect, future studies could add more samples (especially from young and old animals) to the training data.

Our EWAS of age in humpback whale identified 86 CpG sites that changed with age only in females or males, but the vast majority (77%) involved females. Humpback whales are only modestly sexually dimorphic, with females being slightly larger in length than males (Clapham & Mead 1999). However, one difference lies in their investment in reproduction. Females provide all of the maternal investment in offspring after birth. This period of care can last for approximately one year, which includes an extended period of fasting while still on the breeding ground and during migration. Maternal body condition declines during early lactation (Christiansen et al. 2016) and may it account for as much as 41% of caloric intake when feeding resumes (Lockyer 1986; Oftedal 1993). In the Gulf of Maine study population, the reproductive investment of adult females appear to predict their lower average annual apparent survival versus males (Robbins 2007).

We expect that the epigenetic age estimators (epigenetic clock) presented here and by others will have broad applications for study and conservation of humpback whales. There are only a few baleen whale populations where individuals can feasibly be studied from the year of birth. These have provided essential information about age-specific factors, such as age at sexual maturity. However, these likely vary across populations and across time within populations. Furthermore, such data are likely to be biased towards subsets of populations that spend time closer to shore or observers. Unbiased, independent methods of aging are needed that are adequately precise and suitable to living individuals with minimal disturbance. The precision of this clock is particularly important for age class assignment and individual ordination, given that some females produce their first calf as early as five years old (Clapham 1992). Our results further suggests that a clock based on this well-studied baleen whale population may be informative for other populations and species that lack independent age data.

## Methods

### Ethics

Skin samples were collected by the Center for Coastal Studies under research permits issued by the U.S., National Marine Fisheries Service (21485, 16325, 20465, 14245, 633-1483, 633-1778, 932-1905), the Canadian Department of Fisheries and Oceans and IACUC #NWAK-18-02.

### Humpback sample collection

Skin samples were collected from live North Atlantic humpback whales by biopsy sampling techniques (Palsbøll et al. 1991). Sampling was performed in the Gulf of Maine between 2003 and 2020 under the authorization of nationally-issued research permits. Samples were refrigerated or frozen in the field and then archived at −80C without chemical preservative.

Year of birth can be determined precisely for humpback whales first encountered as calves during an obligatory period of maternal dependency (Clapham 1992; Baraff & Weinrich 1993). Otherwise, there are no reliable outward indicators of chronological age in this species. We therefore selected known-age samples for this study based on data from a long-term study of individual whales. The identity of each sampled individual was confirmed through photo-identification techniques (Katona & Whitehead 1981) using a reference catalog of the Gulf of Maine population curated by the Center for Coastal Studies (Provincetown, MA). For individuals first seen as calves, age at sampling was the number of years between sampling and birth. However, the exact date of birth in that year was not known for any individual, and sampling was performed over a wide window (April through November) outside of the winter breeding period. We therefore used an estimated day of birth at the peak of the winter breeding season for all individuals (February 15) to refine age at the time of sampling.

Photo-identification research on this population did not begin until the 1970s and not all of the individuals cataloged since time were first seen as calves. Thus, chronological age is unknown for many individuals, including the oldest whales in the population. We therefore selected additional samples from individuals with long sighting spans (as long or longer than the upper 20% of the available known-age data) to clarify epigenetic age patterns at and beyond the top of the validated age range. In these cases, we calculated a minimum age at sampling based on the fact the individual could not have been born later than the year before their first sighting. However, such whales could have been born in any earlier year and so this provided only a minimum bound on their chronological age.

The sex of sampled individuals was known independently from molecular genetic analysis (Bérubé & Palsbøll 1996), in some cases supplemented by observations of the genital slit (Glockner 1983) or calving history.

### Comparison to Beluga whale skin samples

The skin samples from beluga whales come from a publicly available data set (Bors et al. 2020), which was measured on the same mammalian measurement platform.

### Mammalian methylation array

The mammalian DNA methylation arrays were profiled using the mammalian methylation array (HorvathMammalMethylChip40) (Arneson et al. 2022). The particular subset of species for each probe is provided in the chip manifest file can be found at Gene Expression Omnibus (GEO) at NCBI as platform GPL28271. The SeSaMe normalization method was used to define beta values for each probe (Zhou et al. 2018).

### Penalized Regression models

Details on the clocks (58 CpGs, genome coordinates) and R software code are provided in the Supplementary Table 1 and Supplementary Material. Penalized regression models were created with glmnet (Friedman et al. 2010). We investigated models produced by both “elastic net” regression (alpha=0.5). The optimal penalty parameters in all cases were determined automatically by using a 10 fold internal cross-validation (cv.glmnet) on the training set. The alpha value for the elastic net regression was set to 0.5 (midpoint between Ridge and Lasso type regression) and was not optimized for model performance.

We performed a cross-validation scheme to arrive at unbiased estimates of the accuracy of DNA methylation based age estimators. It consisted of leaving out a single sample (LOOCV) from the regression, predicting an age for that sample, and iterating that procedure across all samples.

### Epigenome wide association studies of age

EWAS was performed for each tissue separately using the R function “standardScreeningNumericTrait” from the “WGCNA” R package (Langfelder & Horvath 2008). Next the results were combined across tissues using Stouffer’s meta-analysis method. The analysis was done using the genomic region of enrichment annotation tool (McLean et al. 2010). The gene level enrichment was done using GREAT analysis and human Hg19 background (McLean et al. 2010).

## Supporting information

Supplementary Table 1. Epigenetic Clocks.

## Acknowledgements

The Center for Coastal Studies conducted the field research to collect tissue samples, as well as well as multi-decade studies of individual humpback whales from their year of birth. Molecular sexing was conducted by the Marine Evolution and Conservation, Groningen Institute for Evolutionary Life Sciences, University of Groningen.

## Data Availability

The data will be made publicly available as part of the data release from the Mammalian Methylation Consortium. Genome annotations of these CpGs can be found on Github https://github.com/shorvath/MammalianMethylationConsortium

The manifest for the Horvath Methylation Array is available at Gene Expression Omnibus (GPL28271: Illumina HorvathMammalianMethylChip40 BeadChip).

