## Supplementary Methods for "DNA methylation age studies of humpback whales"

Technical details surrounding the DNAm age estimator

The epigenetic clock software for humpback whales (and other species) can be applied to data generated on the Mammalian Array Platform (Arneson et al. 2022). New data for epigenetic clock studies can be generated with the mammalian methylation array (HorvathMammalMethylChip40), which is distributed by the Epigenetic Clock Development Foundation: https://clockfoundation.org/

The coefficient values and CpGs underlying the clocks can be found in Supplementary Table 1.

Statistical methods used for building the clocks

The epigenetic clocks were used by employing a single elastic net regression model analysis (R function glmnet). We use used Leave-one-out analysis (LOO) using a single lambda value. We chose the following parameters for the glmnet R function (Alpha: 0.5, CV Fold: 10, Lambda choice for Clock: 1 standard error above minimum CV-MSE).

Covariates and coefficient values of clocks

1)	The humpback whale clock for skin samples is based on 58 CpGs whose coefficient values are specified in the column "HumpbackWhaleSkinClockBasedOnConfidence100". This clock was trained on the basis of animals whose ages were known with 100 percent certainty. We did not use an age transformation.

2)	We also present a second clock for skin samples of humpback whales (55 CpGs) that was trained on the basis of animals whose age was known with at least 90 percent confidence. It specified in the column "HumpbackWhaleSkinClock".

The DNAm Age estimate is estimated as follows. Form a weighted linear combination of the CpGs whose details can be found in the Supplementary Table.

The table reports the probe identifier (cg number) used in the custom Infinium array (HorvathMammalMethylChip40). The weights used in this linear combination are specified in the respective column entitled "Coef.".

The formula assumes that the DNA methylation data measure "beta" values but the formula could be adapted to other ways of generating DNA methylation data.
